# Combinations of antifungals to effectively kill drug-resistant *Candida auris*

**DOI:** 10.1101/776096

**Authors:** Siddharth Jaggavarapu, Eileen M. Burd, David S. Weiss

## Abstract

*Candida auris* is an emerging, highly virulent fungal pathogen that is often resistant to multiple classes of antifungals. To overcome this resistance, we investigated the *in vitro* efficacy of antifungal combinations, finding that amphotericin B/micafungin resulted in drastically reduced survival (~5 logs) of all 10 isolates tested including 2 that were refractory to each individual drug. These findings suggest that combination therapy with micafungin/amphotericin B may have clinical utility against broadly resistant *C. auris*.

## Main Text

Resistance to antifungals is increasing and is particularly concerning in the highly pathogenic *Candida auris*, which is often resistant to multiple classes of antifungal drugs [1]. First discovered in 2009 in the ear canal of a patient in Japan, this fungus has caused outbreaks in multiple healthcare facilities across the world [2-7]. Infections with *C. auris* have high treatment failure rates, resulting in significant mortality [8]. In addition, *C. auris* can reside and survive on contaminated surfaces for up to one week, making eradication difficult [9, 10]. Furthermore, routine clinical diagnostics often misidentify *C. auris* as *Candida haemulonii*, potentially leading to underestimation of its incidence as well as inappropriate treatment [11].

The first line therapy for *C. auris* infections is an echinocandin, with micafungin being the most effective and commonly used [12, 13]. In cases of echinocandin resistance, the polyene amphotericin B is the recommended second line treatment [13]. However, some isolates of *C. auris* are resistant to all three major classes of antifungals; echinocandins, polyenes, and azoles. While research into developing new antifungal drugs that target *C. auris* is ongoing, the limited repertoire of available drugs also necessitates the elucidation of novel methods to more effectively use available antifungals to treat drug-resistant infections [14-16].

We set out to determine the efficacy of combinations of antifungals against 10 isolates of *C. auris* from the FDA/CDC’s Antibiotic Resistance (AR) Bank [17]. Due to the lack of clinical breakpoints for *C. auris*, none of the strains have been classified as resistant, susceptible or intermediate to any of the drugs. A recent study of 123 *C. auris* isolates determined tentative epidemiological cutoff values (ECV), the highest minimum inhibitory concentration (MIC) of the wild-type distribution, for micafungin (0.25 µg/mL), amphotericin B (2 µg/mL) and voriconazole (1 µg/mL) [1]. Therefore, we used >0.25 µg/mL as the putative breakpoint for micafungin, >2 µg/mL for amphotericin B, and >1 µg/mL for voriconazole, respectively. We first tested the efficacy of voriconazole (Acros, Hampton, NH), micafungin (Biovision, Milpitas, CA) and amphotericin B (Millipore, Burlington, MA) individually in preventing the growth of the 10 *C. auris* isolates. Serial dilutions of overnight cultures in YPD broth (Sigma, St. Louis, MO) were plated on YPD agar (Sigma, St. Louis, MO) plates to determine the percentage of surviving, resistant cells. The majority of *C. auris* isolates have been reported to be resistant to azoles [18]. Consistent with that finding, voriconazole was ineffective against all 10 isolates even at 4 µg/mL, resulting in less than a log reduction in survival (Table 1). Testing of micafungin or amphotericin B revealed that 2 isolates were significantly inhibited by either drug, 3 were susceptible to only micafungin, and 3 were susceptible to only amphotericin B. Two strains were refractory to voriconazole, micafungin, and amphotericin B at the concentrations tested (Table 1).

**Table 1.** Survival of *C. auris* isolates plated on antifungal drugs and combinations^*a*^.

| Isolate | no drug | Vori | Mica | Amp B | V+A | V+M | M+A |
| --- | --- | --- | --- | --- | --- | --- | --- |
| 381 | 300,000 | 400,000 | 0 | 0 | 1,000 | 0 | 0 |
| 382 | 700,000 | 700,000 | <10<sup>b</sup> | 0 | 500,000 | 0 | 0 |
| 383 | 1,200,000 | 200,000 | 400,000 | <10<sup>b</sup> | 400,000 | <10<sup>b</sup> | 0 |
| 384 | 600,000 | 600,000 | 600,000 | <10<sup>b</sup> | 500,000 | <10<sup>b</sup> | <10<sup>b</sup> |
| 385 | 300,000 | 200,000 | 200,000 | 200,000 | 600,000 | 10,000 | <10<sup>b</sup> |
| 386 | 800,000 | 1,200,000 | 300,000 | 500,000 | 700,000 | 100,000 | <10<sup>b</sup> |
| 387 | 700,000 | 700,000 | 100,000 | <10<sup>b</sup> | 200,000 | 0 | <10<sup>b</sup> |
| 388 | 500,000 | 300,000 | 2 | 70,000 | 250,000 | 4 | <10<sup>b</sup> |
| 389 | 800,000 | 500,000 | <10<sup>b</sup> | 400,000 | 300,000 | 2 | 2 |
| 390 | 500,000 | 600,000 | 0 | 500,000 | 500,000 | 10 | <10<sup>b</sup> |

<sup>a</sup> Surviving CFUs are shown following plating of the indicated *C. auris* isolates on no drug, voriconazole (Vori, V; 4 µg/mL), micafungin (Mica, M; 0.25 µg/mL), amphotericin B (Amp B, A; 2 µg/mL), or their combinations. Cells within the Table are colored in a gradient from highest CFUs (red) to lowest (green).
<sup>b</sup> CFUs could not be determined exactly since dense growth was observed at the highest concentration of bacteria, but no growth was observed at the next dilution.

The combination of voriconazole and amphotericin B was ineffective at killing any of the strains. In fact, a few isolates that were completely inhibited by amphotericin B, survived better when treated with the combination of voriconazole and amphotericin B, indicative of antagonism (Table 1). Indeed, prior studies have concluded that some azoles and amphotericin B can be antagonistic since these azoles inhibit the biosynthesis of ergosterol, the target of amphotericin B [19].

The combination of voriconazole and micafungin resulted in a ~5 log reduction in survival of 8 of the 10 isolates (Table 1). Importantly, 3 isolates resisted voriconazole and micafungin individually, but were inhibited by their combination, suggesting a synergistic effect of these two antifungals. However, the two isolates that were resistant to the individual effects of voriconazole, micafungin, and amphotericin B, were also uninhibited by voriconazole/micafungin.

Importantly, the combination of micafungin and amphotericin B resulted in a ~5 log reduction in survival of all 10 isolates (Table 1). Notably, this combination was effective against strains 385 and 386 that we classified as resistant to all three individual antifungals tested (Table 1). These results suggest that micafungin/amphotericin B may be able to overcome broadly resistant *C. auris* isolates and offer an effective treatment option using existing drugs and without the need for development of novel antifungals.

Since the current recommended first line treatment for *C. auris* infections is an echinocandin (e.g. micafungin), we tested if initial exposure to micafungin would alter the subsequent efficacy of the micafungin/amphotericin B combination, or the current recommended second line treatment of amphotericin B alone. We grew the 5 isolates that were resistant to micafungin overnight in YPD broth containing a sublethal concentration (0.0625 µg/mL) of the drug and subsequently plated onto YPD agar containing micafungin/amphotericin B or amphotericin B alone, to quantify survival. Isolates 383, 384 and 387 that were responsive to amphotericin B monotherapy (Table 1) prior to micafungin treatment were still susceptible to amphotericin B after exposure to sublethal micafungin, suggesting that prior exposure to micafungin did not affect susceptibility to amphotericin B (Table 2). Interestingly, the combination of micafungin/amphotericin B was also still effective against micafungin-pretreated isolates (Table 2). These results suggest that combination therapy with micafungin/amphotericin B may be effective as a second line regimen even after failed treatment with micafungin monotherapy.

**Table 2.** Survival of *C. auris* after initial exposure to sublethal micafungin^*a*^.

| Isolate | no AB | Mica | AmpB | M+A |
| --- | --- | --- | --- | --- |
| 383 | 100,000 | 200,000 | 10 | 0 |
| 384 | 60,000 | 60,000 | 0 | 0 |
| 385 | 40,000 | 20,000 | 2,100 | 0 |
| 386 | 80,000 | 20,000 | 20,000 | 0 |
| 387 | 40,000 | 10,000 | 0 | 0 |

<sup>a</sup> All isolates were initially grown overnight in the presence of 0.0625 µg/mL micafungin. Subsequently, isolates were plated onto no drug (“no AB”) or the indicated drugs at concentrations of 0.25 µg/mL micafungin (Mica, M) and/or 2 µg/mL amphotericin B (AmpB, A). Cells within the Table are colored in a gradient from the highest CFUs (red) to the lowest (green).

The results presented here reveal that the combination of micafungin and amphotericin B has broad efficacy against a collection of *C. auris* clinical isolates, including several strains that we classified as exhibiting resistance to all 3 major antifungal classes. This regimen may have enhanced efficacy as compared to amphotericin B, the current second line therapy for echinocandin resistant strains, and may justify a re-examination of the treatment guidelines. While the results are encouraging, further testing of this combination regimen against clinical isolates will be critical. If these results are representative and broadly applicable, they suggest that the combination of micafungin/amphotericin B may offer an effective option to treat infections caused by highly resistant *C. auris*.

